# Pan-Cancer Repository of Validated Natural and Cryptic mRNA Splicing Mutations

**DOI:** 10.1101/474452

**Authors:** Ben C. Shirley, Eliseos J. Mucaki, Peter K. Rogan

## Abstract

We present a major public resource of mRNA splicing mutations validated according to multiple lines of evidence of abnormal gene expression. Likely mutations present in all tumor types reported in the Cancer Genome Atlas (TCGA) were identified based on the comparative strengths of splice sites in tumor versus normal genomes and then validated by respectively comparing counts of splice junction spanning and abundance of transcript reads in RNA-Seq data from matched tissues and tumors lacking these mutations. The comprehensive resource features 351,423 of these validated mutations, the majority of which (69.1%) are not featured in the Single Nucleotide Polymorphism Database (dbSNP 150). There are 117,951 unique mutations which weaken or abolish natural splice sites, and 244,415 mutations which strengthen cryptic splice sites (10,943 affect both simultaneously). 27,803 novel or rare flagged variants (with <1% population frequency in dbSNP) were observed in multiple tumor tissue types. Single variants or chromosome ranges can be queried using a Global Alliance for Genomics and Health (GA4GH)-compliant web Beacon, Validated Splicing Mutations, either separately or in aggregate alongside other beacons through the public Beacon Network (http://www.beacon-network.org/#/search?beacon=cytognomix), as well as through our website (https://validsplicemut.cytognomix.com/).

## Introduction

Next generation sequencing continues to reveal large numbers of novel variants whose impact cannot be interpreted from curated variant databases or through reviews of peer-reviewed biomedical literature^1^. This has created a largely, unmet need for unequivocal sources of information regarding the molecular phenotypes and potential pathology of variants of unknown significance (VUS); in cancer genomes, such sources are critically needed to assist in distinguishing driver mutations from overwhelming numbers of bystander mutations. VUS classification criteria highlight the limitations in genome interpretation due to ambiguous variant interpretation. Of the 458,899 variant submissions in NCBI’s ClinVar database with clinical interpretations, nearly half (n=221,271) are VUS (as of November 5th 2018; https://www.ncbi.nlm.nih.gov/clinvar/submitters/). Only 10,784 variants in ClinVar have been documented to affect mRNA splicing at splice donor or acceptor sites, with 1,063 of these being classified as VUS, and cryptic mRNA splicing mutations are not explicitly described. The current ACMG criteria^2^ for variant pathogenicity prevent clinical classification of most VUS. Functional evidence that VUS either disrupt or abolish expression of genes has been sought to improve classification and provide insight into the roles, if any, of individual VUS in predisposing or causing disease. We present a comprehensive data repository for a relatively common mutation type (cis-acting variants that alter mRNA splicing). Mutations are predicted with information theory-based analyses^3^, and supported with functional evidence that variants in tumor genomes are specifically associated with abnormally spliced mRNAs that are infrequent or absent in transciptomes lacking these variants^4^.

Information Theory (IT) has been proven to accurately predict impact of mutations on mRNA splicing, and has been used to interpret coding and non-coding mutations that alter mRNA splicing in both common and rare diseases^3,5–15^. We have described an IT-based framework for the interpretation and prioritization of non-coding variants of uncertain significance, which has been validated in multiple studies involving novel variants in patients with history or predisposition to heritable breast and/or ovarian cancer ^11–15^.

The Cancer Genome Atlas (TCGA) Pan-Cancer Atlas (PCA) is a comprehensive integrated genomic and transcriptomic resource containing data from >10,000 tumors across 33 different tumor types^16^. Here, we utilized IT-based tools for assessment of high quality sequenced variants in TCGA patients for their potential impact on mRNA splicing. The accuracy of predicted mutations was evaluated with an algorithm we previously developed that compares transcripts from individuals carrying these variants with others lacking them. The results of these genome-wide analyses are presented using an online resource (https://validsplicemut.cytognomix.com/) which can be queried through the Beacon Network (https://beacon-network.org)^17^.

## Materials & Methods

### TCGA Data Acquisition and Processing

Controlled-access data was obtained with permission from the Data Access Committee at NIH for TCGA and from the International Cancer Genomics Consortium. Patient RNA sequencing BAM files (tumor and normal, when available) and their associated VCF files (HG19) were initially obtained from the CancerGenomeHub (CGhub). Files were later downloaded through Genomic Data Commons (https://gdc.cancer.gov/) using the GDC Data Transfer Tool, as CGhub was decommissioned mid-project. Variants in VCF files which did not pass quality control (QC) were not analyzed.

### Information Analysis and RNA-Seq Validation of Splicing Variants

We used the *Shannon Pipeline* software (which applies IT to rapidly perform high-throughput, *in silico* prediction of the impacts of variants on mRNA splicing)^18^ to analyze all QC-passing variants in VCFs from TCGA (>168 million variants) to evaluate their potential impact on splice site binding strength (changes in information content, R_*i*_, measured in bits). Variants which were predicted to strengthen known natural sites or weaken cryptic splice sites were excluded from all subsequent analyses.

To validate the potential impact of Shannon Pipeline-flagged mutations, *Veridical* software analyzed genomic variants (including insertions and deletions) by comparing the RNA-Seq alignment in the region surrounding the variant with the corresponding interval in control transcriptomes (normal and tumor tissue of the same type) lacking the variant^4,19^. Veridical: a) counts abnormally spliced reads in RNA-Seq data (categorized as: cryptic site use, exon skipping, or intron inclusion [containing or adjacent to the flagged mutation]), b) applies the Yeo-Johnson transformation to these results, and c) determines the null hypothesis probability (p-value) that the transformed read count corresponds to normal splicing. In tumor types where normal controls were not available, a set of RNA-Seq datasets from 100 different normal tissues from TCGA were used (e.g. a combination of 5 tissue types: BRCA, BLCA, LUAD, KIRC, PRAD). Veridical results that were not significant for a particular variant (p-value > 0.05 for all of the splicing categories) were not further analyzed. After analysis, Veridical validated 351,423 unique mutations for their direct impact on mRNA splicing (Table 1). The Shannon pipeline-flagged and Veridical-filtered results were combined into a single large table (*Dataset 1*), the source data for the ValidSpliceMut SQL database and the associated Beacon application.

**Table 1:**
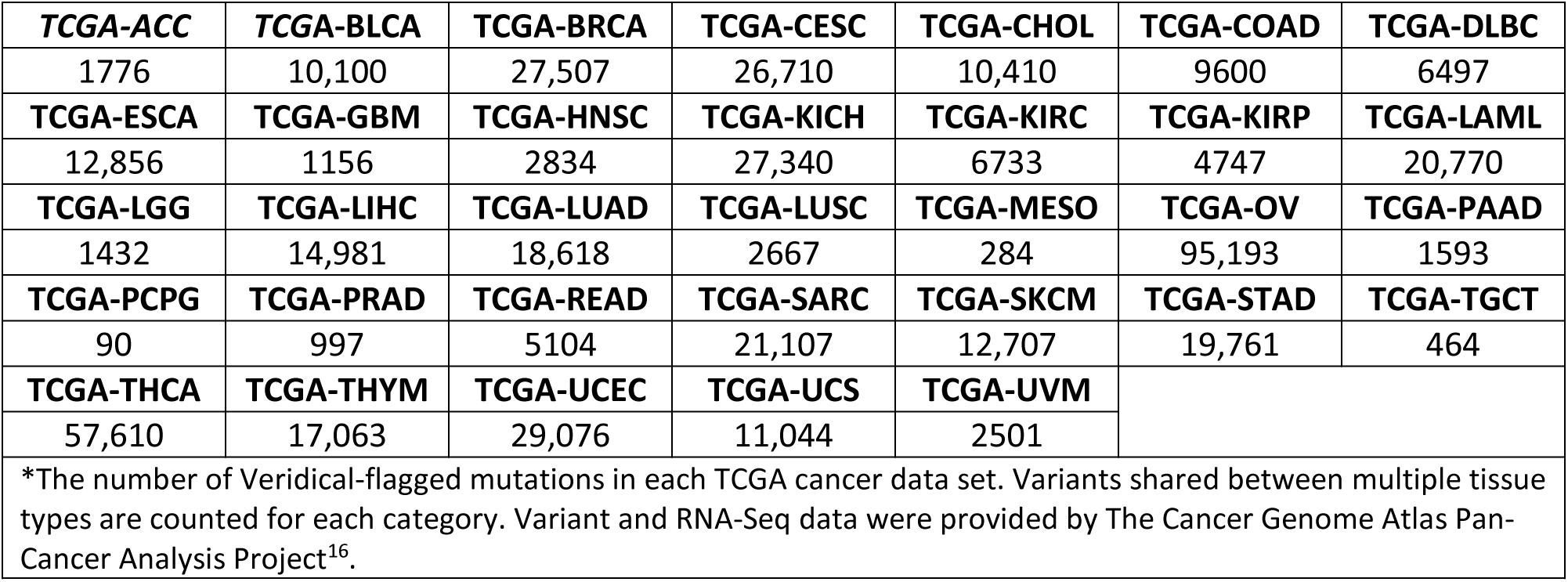
Unique Flagged Variants by TCGA Tumor Tissue Type^*^

### Development of the ValidSpliceMut database and Beacon

We created a publicly accessible Application Programming Interface (API) (https://beacon.cytognomix.com) that can be utilized to programmatically query variants passing filter thresholds described above (*Dataset 1*). It was built in accordance with the GA4GH Beacon v1.0.0 specification (https://github.com/ga4gh-beacon/specification/blob/master/beacon.md), which describes a Representational State Transfer (REST) API for genetic data sharing. A beacon accepts queries using an HTTP request and returns JavaScript Object Notation (JSON) object. Our Beacon implementation is coded in PHP 7.0 and utilizes a MySQL database with indexes applied to variant ID, chromosome, and coordinate fields. The returned JSON object reports whether the variant was found within our Beacon dataset as well as metadata including splice site coordinate, splice type, site type, the IT-based measures *R*_*i,initial*_, *R*_*i,final*_, affected individual IDs, tumor type, Veridical evidence by type annotated with significance level, and, if known, the corresponding rsID with its average heterozygosity (dbSNP 150). The metadata for each variant sent the beacon network is a concise subset of available results in our database. It includes the first relevant database entry, meaning that if the variant exists within multiple individuals only the first will contribute fields to the metadata. However, among this metadata is a hyperlink to our local website containing results for any remaining tumors.

We developed the website ValidSpliceMut (https://validsplicemut.cytognomix.com; Figure 1) to serve as a local interface to our beacon, allowing users to manually input a variant or coordinate range, automatically query the API with the request, and view formatted results. This website provides a complete view of variants, including Veridical-based evidence on all data related to every affected individual. If a variant is associated with multiple splice sites, the user is presented with a brief overview of all affected sites and must select a desired site to continue. To obtain the coordinate of the queried variant in gene-centric notation, a link is provided which queries the Mutalyzer API (https://mutalyzer.nl/json/) and generates coordinates for all available transcripts. ValidSpliceMut only reports transcripts for the gene affected by the variant.

**Figure 1:**
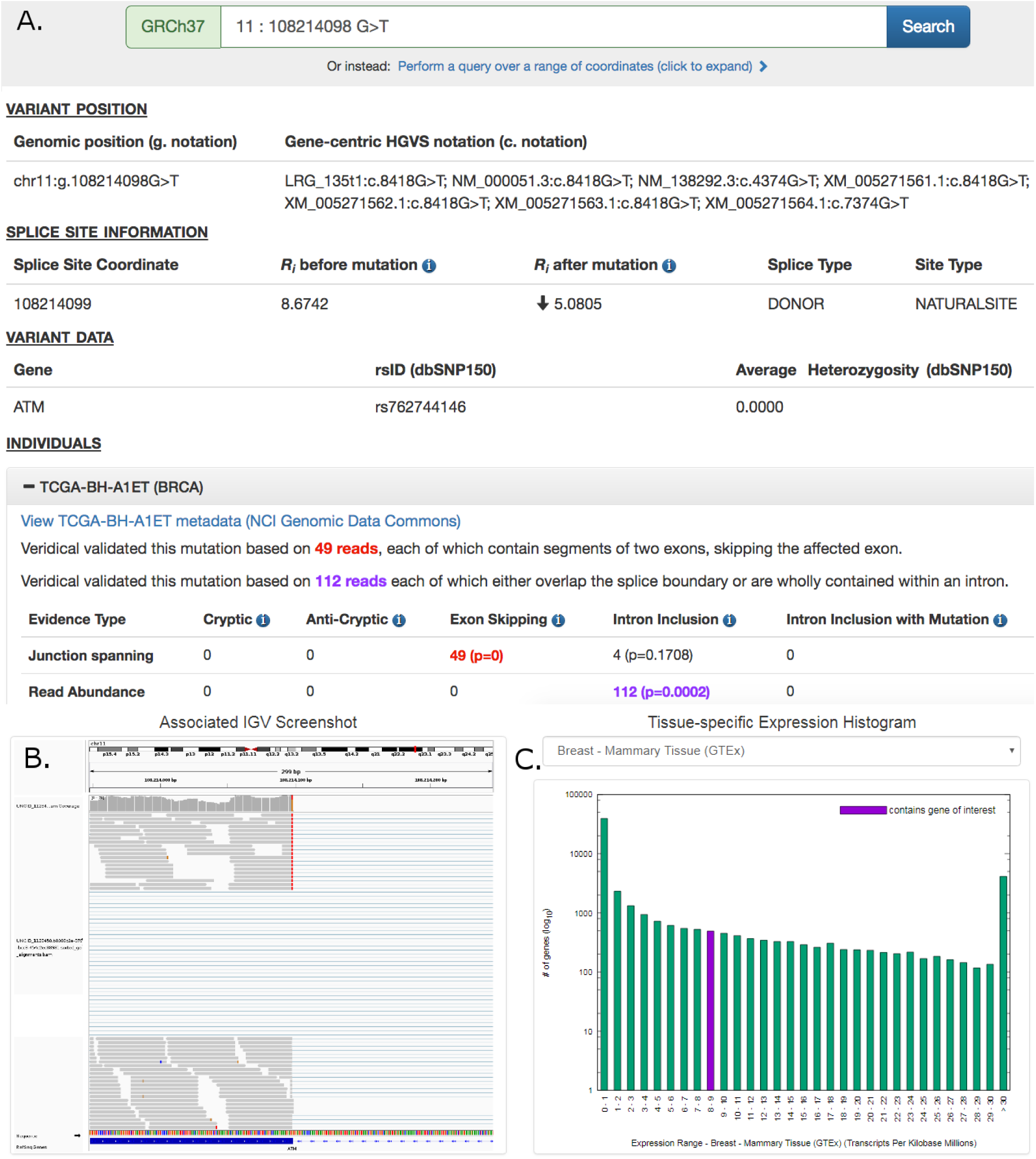
Screenshot of *ATM*:g.108214098G>T Results Provided By ValidSpliceMut Website. (A) The ‘Variant Position’ heading displays the variant of interest in g. notation, and provides a link which queries the Mutalyzer API to obtain the variant coordinate in a gene-centric c. mutation format. Variant-specific and splice site-specific tabular results are presented under the headings “Splice Site Information” and “Variant Data”. Results are organized by TCGA sample IDs harboring the mutation within a series of expandable panels. A link is provided to patient tumor metadata on the GDC data portal. Each panel consists of read counts and p-values by Veridical evidence type. Significant p-values (< 0.05) are highlighted in bold. Evidence types deemed “strongly corroborating” (Viner et al. 2014) are color coded and correspond to the dynamically generated text appearing above the table. (B) An IGV image showing alignment of expressed sequence reads. IGV screenshots are provided only for mutations present <1% of population (in dbSNP 150), with ≥ 5 junction-spanning reads, and are highly significant (p < 0.01) for cryptic splicing, exon skipping, and/or intron inclusion with mutation. A specific IGV screenshot for this sample captures the region surrounding the mutation. Here, several RNA-Seq reads show skipping of the affected exon. (C) A dynamically generated histogram presents expression levels of all genes for a selected normal tissue type. Genes are grouped into bins based on expression level, denoted on the x-axis. The number of genes present in each bin is shown on the y-axis (log_10_ scale). The histogram key indicates the expression range of the variant-containing gene. Tissue type can be changed via a drop-down list.

A Results page presents variant-specific data in tabular format and an expandable list of panels describing the affected individuals. Each of these panels contains Veridical output in tabular format for the selected tumor, a link to the tumor metadata at US National Cancer Institute (by querying the GDC API [https://api.gdc.cancer.gov/] to obtain a UUID which is used to construct a link to the GDC data portal), an Integrative Genome Viewer (IGV) screenshot containing the variant (IGV screenshots are available for selected variants, see below), and a histogram which presents the expression levels of the variant-containing gene compared to all other gene expression levels across a selected normal tissue type (created dynamically using gnuplot 5.0). The tissue expression data is provided by GTEx (https://gtexportal.org/home/; downloaded on 10/22/18). However, several TCGA tumor types did not have a GTEx equivalent (CHOL, DLBC, MESO, READ, SARC, THYM and UVM). The GNF Expression Atlas 2 ^20^ was used for expression data for both lymph nodes (DLBC) and the thymus (THYM). For the remaining tissues, expression data from the following studies were obtained from the e Genome Expression Omnibus (GEO): <u>GSE76297</u> (CHOL), <u>GSE2549</u> (MESO), <u>GSE15781</u> (READ) <u>GSE44426</u> (SARC), and <u>GSE44295</u> (UVM).

To generate IGV images presented on the webpage, a bash script was written to automatically load the RNA-Seq BAM file of a patient with a mutation of interest into IGV, set the viewing window within the region of interest (300nt window, centered on the variant), sorted to bring reads containing the variant of interest to the top of the screen (to increase chance of visualizing mutant splice form), followed by a screen capture. The generation and storage of IGV images for all patient-mutation pairs would be prohibitive due to limitations in time and server space requirements. Therefore IGV images showing evidence of splicing abnormalities were generated *only* for patient-mutation pairs which met the most stringent criteria: the mutation was required to be flagged for junction-spanning cryptic site use, exon skipping, or intron inclusion (with mutation); the flagged category must include 5 or more reads in this category; if the variant is present in the dbSNP database (release 150), the frequency was required to be < 1% of the population; and the Veridical results, in which the mutations flagged were required to exhibit p ≤ 0.01 for at least one form of evidence of a splicing abnormality. In some cases, the splicing event observed by Veridical may not be present within the image window as the automated procedure used to create these images does not present all evidential sequence reads due to limitations on the number of reads that are shown. Additionally, reads appearing as exon skipping may instead indicate a pre-existing cryptic site outside of the viewing window (see Table 2; *FAT1*:g.187521515C>A [c.11641-1G>T] and *SMAD3*:g.67482748C>G [c.1155-3C>G]).

**Table 2:**
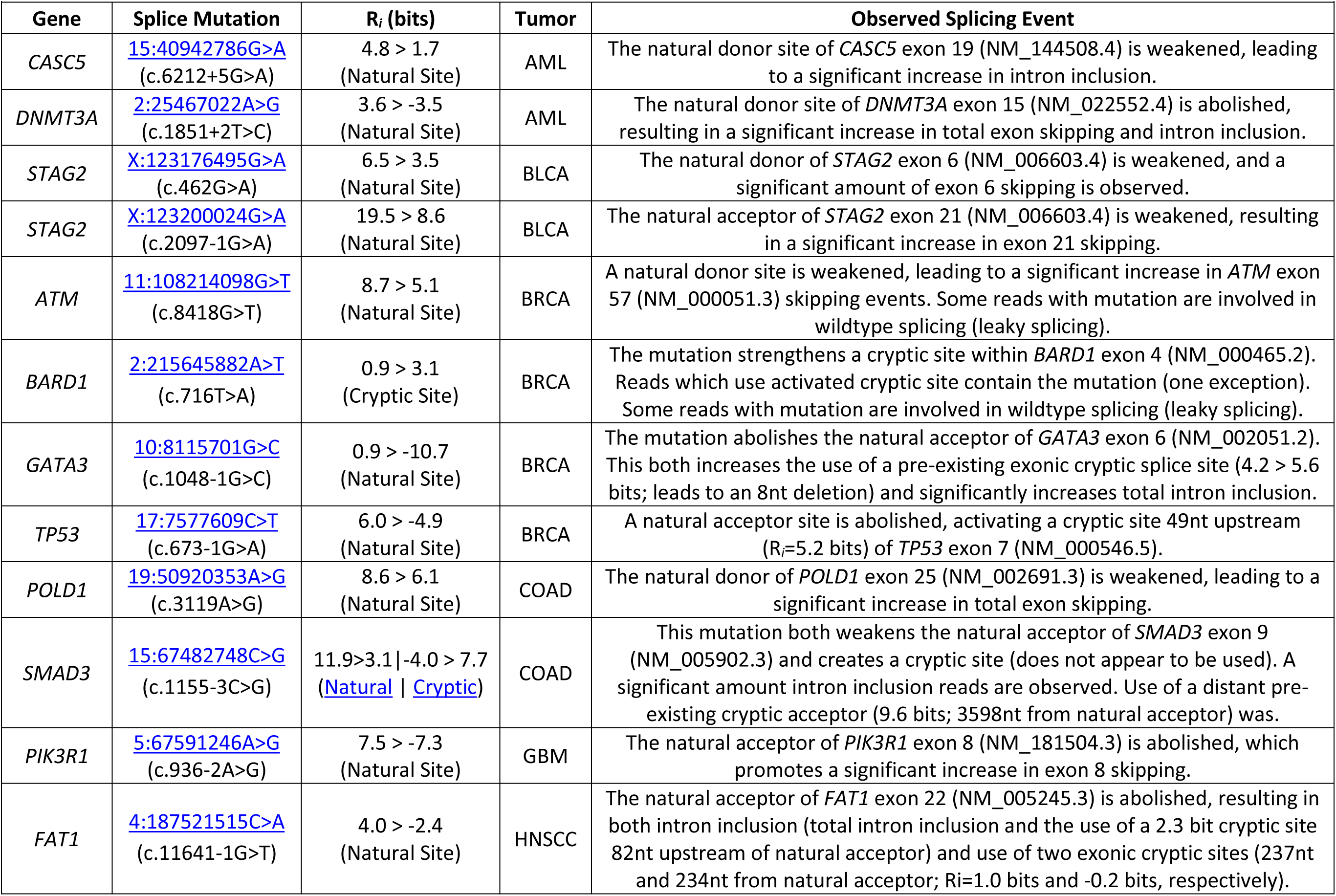
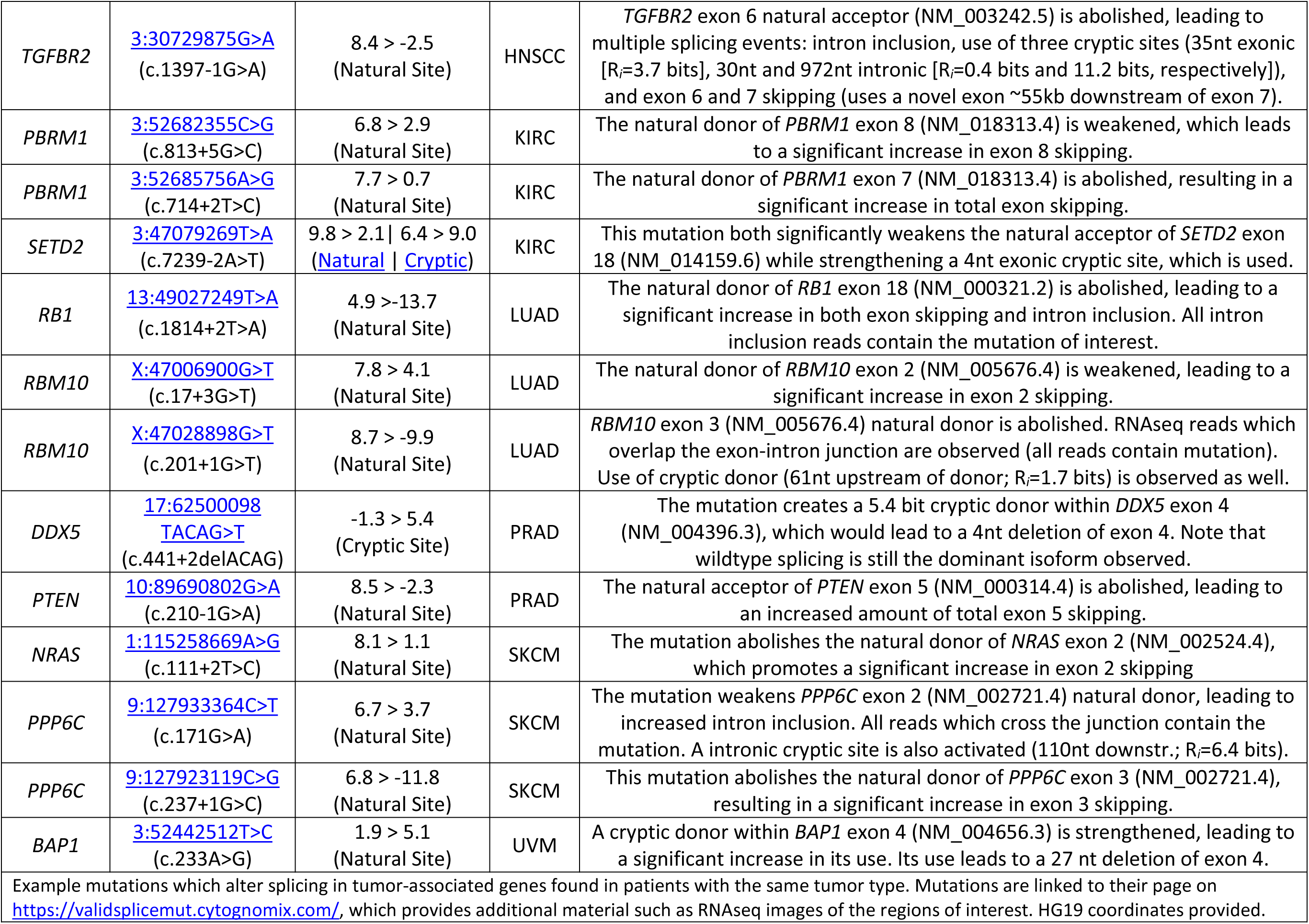
Validated Splicing Mutations in COSMIC Cancer Gene Census genes in TCGA tumor genomes

## Results and Discussion

We have derived a GA4GH-standardized, searchable web resource for a large set of validated mRNA splicing variants present in diverse tumor types. All variants passing QC in TCGA cancer patients were analyzed with the Shannon pipeline^18^. This revealed that 1,297,242 variants were predicted to have significant impacts on normal mRNA splicing (347,549 natural and 985,112 cryptic splice sites; 35,419 affecting both types). Subsequent RNA-Seq analysis with Veridical^4^ provided evidence of abnormal gene expression specifically associated with a subset of these variant(s), identifying 351,423 unique mutations. Results are searchable through either the Beacon Network (https://beacon-network.org), or our publicly-accessible webpage (https://validsplicemut.cytognomix.com/).

Our results contrast with another TCGA study that investigated alternative mRNA splicing^21^ and demonstrated a limited set of non-constitutive exon-exon junctions attributable to cis-acting splicing mutations (n = 32). The 2,736 novel or rare variants that we report which specifically activate cryptic splicing (significant ‘junction-spanning cryptic site use’ reads found by Veridical), exceed the number reported in another study that analyzed all available TCGA tumor transcriptomes (n=1,964)^22^.

Validated variants were also tallied by tumor tissue type in our study (Table 1). 33.6% of unique mutations (n=117,951) significantly weaken natural splice sites, while 69.6% (n=244,415) strengthen novel or pre-existing cryptic sites. 242,983 variants (69% of all flagged variants) are absent from dbSNP150. 73,975 variants (21%) are found in <1% of the population. Valid mutations lacking rsIDs represent either novel or recently observed variants. This low level of dbSNP saturation is consistent with the idea that many currently unknown mRNA splicing mutations may yet be discovered through additional sequencing studies.

In Table 2, we highlight a subset of validated splicing mutations (n=25) which were identified in known driver genes implicated in the COSMIC (Catalogue Of Somatic Mutations In Cancer; https://cancer.sanger.ac.uk/cosmic) Cancer Gene Census catalog (CGC)^23^. These mutations are associated with either increased exon skipping, intron inclusion, and/or cryptic site use. Mutations in Table 2 are hyperlinked to the ValidSpliceMut webpage which provides additional information, including expression evidence supporting predictions made by the Shannon pipeline.

Many mutations generated multiple types of abnormal read evidence present in misspliced transcripts. Interestingly, a subset of mutations (n=28) produced evidence for every type of abnormal splicing reported by Veridical. *Dataset 2* (see Data Availability) describes 11 representative mutations that simultaneously increase exon skipping, intron inclusion, and activate (or significantly increase utilization of) a strengthened cryptic site. In all but one instance, the mutation weakens the natural site while simultaneously strengthening a nearby cryptic site. The one exception involves the gene *SAP30BP*, where simultaneously occurring mutations in the same read (in linkage disequilibrium; separated by 4 nucleotides) independently cause two separate splicing changes: g.73702087G>A (c.661-1G>A; abolishes the natural acceptor of exon 10) and g.73702091G>A (c.664G>A; creates a weak cryptic acceptor site). The combined splicing impact of these variants is significant exon skipping, intron inclusion, and use of the activated cryptic site.

Because of the requirement for expression validation, this resource presents a set of splicing abnormalities in which we have the highest confidence. We anticipate that some correct predictions of the Shannon pipeline may have not been validated by Veridical due to the limitations of mRNA detection; for example, either low expression of the gene harboring the mutation or nonsense-mediated decay of the corresponding transcript could be consistent with the effects of a valid splicing mutation, but in the absence of a sufficient number of abnormal reads, the mutation could not be confirmed. Furthermore, at the time that the current analysis was performed, the available Shannon pipeline version did not report regulatory splicing variants adjacent to constitutive and cryptic splice sites which influence exon definition. Due to the substantial processing required for the complete TCGA dataset, the present analysis does not incorporate the effects of these variants on exon definition, which we have modeled by IT^6^; it does not predict the relative abundance of leaky, natural and cryptic isoforms, though such information might be inferred from the expression data on each tumor. The current version of Shannon pipeline does integrate predictions of splicing regulatory sequences and accounts for relative abundance of mRNA isoforms by exon definition, and is available through the MutationForecaster system (https://mutationforecaster.com).

The Validated Splicing Mutation resource should substantially contribute to reducing the number of outstanding VUS in tumor (and possibly some germline) genomes, and substantially increases the number of splicing-related variants based on previously unappreciated molecular consequences, in particular, activation of cryptic splice sites. In our previous study^19^, a subset of the TCGA breast cancer patient data was evaluated with IT-based tools, identifying 988 variants as significantly altering normal splicing by Veridical (19% of total mutations flagged by IT). This database greatly expands the size of the repository. Here, a higher ratio of rare or novel mutations have been validated by Veridical (24% of total mutations were flagged by IT). The higher yield found could be related to the same mutation being present in multiple samples from the same tumor type and other tumor tissues, which would be expected to increase the probability of observing abnormally expressed splice forms for the mutation.

### Data Availability

#### *Zenodo*: Dataset 1. Validated natural and cryptic mRNA splicing mutations

Source data computed by the Shannon pipeline and Veridical, displayed on the ValidSpliceMut website (https://validsplicemut.cytognomix.com/). DOI: <u>10.5281/zenodo.1488211</u>

#### *Zenodo:* Dataset 2. Mutations which lead to multiple types of aberrant splicing

Representative set of mutations which significantly alter splicing in all evidence types analyzed by Veridical (i.e. cryptic splice site use, exon skipping, intron inclusion). Mutations are linked to their page on https://validsplicemut.cytognomix.com/, which provides additional material such as RNA-Seq images of the regions of interest. DOI: <u>10.5281/zenodo.1489941</u>

License: <u>CC0 1.0</u>

#### Consent

Controlled-access TCGA sequence data was accessed with permission from NCBI (dbGaP Project #988: “Predicting common genetic variants that alter the splicing of human gene transcripts”; Approval Number #13930-11; PI: PK Rogan) and the International Cancer Genome Consortium (ICGC Project #DACO-1056047; “Validation of mutations that alter gene expression”).

#### Author Contributions

PKR designed the methodology, obtained approved access to the TCGA data and oversaw the project. EJM downloaded and processed the data, and performed analyses on said data. BCS designed and built the Beacon software and the ValidSpliceMut webpage. BCS, EJM and PKR wrote the manuscript.

## Acknowledgements

We acknowledge Coby Viner, Stephanie Dorman, Will Phillips and Ujani Hazra for their contributions to the early stages of this project. We are grateful to Max Barkley, Milan Panik and Miro Cupak (DNAStack) for their assistance in integrating our ValidSpliceMut beacon into the GA4GH network.

